# Kin recognition in guppies uses self-referencing on olfactory cues

**DOI:** 10.1101/2020.05.28.122275

**Authors:** Mitchel J. Daniel, F. Helen Rodd

**Affiliations:** Department of Ecology and Evolutionary Biology, University of Toronto, Toronto, Ontario, CA; Department of Biological Science, Florida State University, Tallahassee, Florida, USA

**Author notes:** **Corresponding Author**, Mitchel J. Daniel, 319 Stadium Drive, FL 32304, Florida State University, Tallahassee, FL, USA.

**Keywords:** Kin recognition, Inbreeding avoidance, Male-male competition, Nepotism, Trinidadian guppy (*Poecilia reticulata*), Olfaction, Multiple paternity

## Abstract

Kin recognition plays an important role in social evolution, but the proximate mechanisms by which individuals recognize kin remain poorly understood. In many species, individuals form a “kin template” that they compare against conspecifics’ phenotypes to assess phenotypic similarity–and by association, relatedness. Individuals may form a kin template through self-inspection (i.e. self-referencing) and/or by observing their rearing associates (i.e. family-referencing). However, despite much interest, few empirical studies have successfully disentangled self- and family-referencing. Here, we use a novel set of breeding crosses in the Trinidadian guppy (*Poecilia reticulata*) to definitively disentangle referencing systems by manipulating exposure to kin from conception onwards. We show that guppies discriminate among their full- and maternal half-siblings, which can only be explained by self-referencing. Additional behavioral experiments revealed no evidence that guppies incorporate the phenotypes of their broodmates or mother into the kin template. Finally, by manipulating the format of our behavioral tests, we show that olfactory communication is both necessary and sufficient for kin discrimination. These results demonstrate that individuals recognize kin by comparing the olfactory phenotypes of conspecifics against their own. This study resolves key questions about the proximate mechanisms underpinning kin recognition, with implications for the ontogeny and evolution of social behavior.

## Introduction

Social interactions are nearly ubiquitous in nature, and can have profound consequences for fitness. Kin recognition systems allow individuals to assess their relatedness to conspecifics and adjust their behavior accordingly, facilitating behavioral strategies that increase the fitness consequences of their social actions. Such strategies include inbreeding avoidance, nepotism, and the formation of cooperative kin groups (reviewed in Hauber et al. 2001; Penn and Frommen 2010). Evidence of kin recognition has been broadly reported across animal taxa (Penn and Frommen 2010); however, fundamental aspects of the proximate mechanisms that underpin kin recognition remain poorly understood.

Many species rely on a kin recognition system called phenotype matching, wherein individuals use phenotypic similarity as a signal of relatedness (Hauber and Sherman 2001; Mateo and Holmes 2004; Penn and Frommen 2010; Krupp and Taylor 2013). To do so, the evaluating individual first acquires information about its own phenotype through self-inspection (i.e. self-referencing) and/or by observing the phenotypes of the putative kin with which it develops, such as its broodmates or parents (i.e. family-referencing). The evaluator stores this phenotypic information internally as a so-called “kin template”. Subsequently, the evaluator can compare the kin template against the phenotype of a given conspecific, and infer relatedness from the degree of similarity between the two.

Phenotype matching is a versatile kin recognition system because it can be used to recognize familiar and unfamiliar kin alike (Mateo and Holmes 2004; Penn and Frommen 2010). In principle, phenotype matching could rely on any phenotypic cue(s), so long as there is a robust association between phenotypic and genotypic similarity (Penn and Frommen 2010; Krupp and Taylor 2013). However, for many species, it remains unclear what cues are used, and whether the kin template is acquired through self- or family-referencing (Hauber and Sherman 2001; Tang-Martinez 2001; Penn and Frommen 2010).

Whether phenotype matching is self- or family-referential can have important implications for the ontogeny and evolution of phenotype matching. These two referencing systems invoke different cognitive and sensory processes (Hauber and Sherman 2001; Mateo and Holmes 2004), and their relative reliability should differ depending on the natural history of the organism. For example, family-referencing is a more direct means of recognizing broodmates and/or parents, and allows the use of phenotypes that cannot be readily assessed through self-inspection (e.g. some visual cues). However, accurate family-referencing requires that early life social experience be limited to close relatives; otherwise, the phenotypes of non-kin could be integrated into the kin template (Hauber and Sherman 2001; Penn and Frommen 2010). When early life social experience is not limited to close relatives, self-referencing is expected to evolve instead (Holmes and Sherman 1982; Hauber and Sherman 2001). For example, high levels of multiple paternity reduce relatedness amongst broodmates, diminishing the accuracy of a family-referent kin template, and are therefore predicted to select for self-referencing (Hauber and Sherman 2001; Hain and Neff 2006).

Experimentally distinguishing between self- and family-referencing has proven challenging, because its requires manipulation of the phenotypic cues that individuals are exposed to throughout ontogeny without disrupting natural development (Hauber and Sherman 2001). Numerous attempts have relied on cross-fostering or socially isolating individuals shortly after birth, or as eggs (e.g. Blaustein and O’Hara 1981, 1983; Buckle and Greenberg 1981; Holmes and Sherman 1982; Getz and Smith 1986; Holmes 1986; Aldhous 1989; Smith et al. 1994; Todrank et al. 1998; Mateo and Johnston 2000; Hesse et al. 2012). The reasoning behind these approaches is that, if an individual develops apart from its true kin, only self-referencing should lead to an accurate kin template; therefore, the presence or absence of accurate kin recognition implicates self- or family-referencing, respectively (Mateo and Holmes 2004). These approaches have yielded useful insight into the ontogeny of kin recognition, but have also been criticized because they may not fully eliminate exposure to the cues of broodmates or the mother (Hare et al. 2003; Mateo and Holmes 2004; Robinson and Smotherman 2005). In zebrafish, family-referencing requires only 24 h exposure to kin (Gerlach et al. 2008). Thus, even very brief exposure to kin between birth and separation could be sufficient for the development of family-referencing. Furthermore, developing embryos may be exposed to the cues of kin while in the egg (e.g. Hauber et al. 2001) or, for live-bearing species, *in utero*. The uterine environment can shape sensory system development and behavioral preferences later in life. For example, odor cues in the amniotic fluid can enhance subsequent detection of, and preference for, those odors (Bellinger et al. 2004; Todrank et al. 2011). *In utero* exposure to biologically important compounds, which can include metabolites and protein products of the mother or broodmates (Bellinger et al. 2004), may therefore facilitate formation of a family-referent kin template during gestation (Robinson and Smotherman 2005; Penn and Frommen 2010). However, investigations into this possibility remain lacking. Furthermore, few studies have satisfactorily disentangled self- and family-referencing by accounting for potential exposure to the cues of kin throughout all of development, from conception onwards (but see Hain and Neff 2006).

Here, we report an experiment that distinguishes between self- and family-referencing in the Trinidadian guppy (*Poecilia reticulata*). In this live-bearing fish, phenotype matching facilitates female inbreeding avoidance (Daniel and Rodd 2016), male avoidance of intrasexual competition with kin (Daniel and Williamson 2020), and, in some populations, preferential shoaling with close relatives by juveniles (Griffiths and Magurran 1999; Hain and Neff 2007). However, whether guppies use self- or family-referencing is, to our knowledge, untested. *In utero* exposure to the cues of the mother or broodmates could provide an opportunity for the development of family-referencing in the guppy, though the high rates of multiple paternity in many guppy populations should favour the evolution of self-referencing (Hain and Neff 2007; Johnson et al. 2010). While little is known about the sensory system development of guppy embryos, other teleost fish such as zebrafish express olfactory receptors that become sensitive to odorants well before embryonic development is completed (Barth et al. 1996; Li 2005). Additionally, guppies in some wild populations form size-assortative shoals shortly after birth that may consist primarily of broodmates (Piyapong et al. 2011). Thus, there may be ample opportunities for self-referencing and family-referencing *in utero* and after birth. Rather than use traditional methods of cross-fostering or social isolation that would not entirely disentangle these mechanisms, we employ a combination of breeding crosses that provide a novel way to distinguish self- and family-referencing while preserving natural development.

We also investigate the cues that guppies use in phenotype matching, by determining the sensory modality of kin recognition. Both visual and olfactory cues are known to influence mate choice (e.g. Houde 1987, 1997*a*; Shohet and Watt 2004) and shoaling partner choice (Houde 1997; Griffiths and Magurran 1999; Shohet and Watt 2004) in this species, suggesting that both modalities could be important for phenotype matching. Here, we determine the relative importance of each modality by assaying kin discrimination in formats that allowed only visual, only olfactory, or both visual and olfactory communication.

## Methods

### Overview of experiments

We investigated the proximate mechanisms that underlie phenotype matching in guppies using a series of four behavioral experiments. First, we tested whether guppies use self-referencing to recognize kin by asking whether they discriminate between their full-siblings and maternal half-siblings, both of which they developed with *in utero* and after birth. Second, we tested whether guppies use family-referencing based on the phenotypes of their broodmates. We asked this by determining whether guppies discriminate between two non-relatives, when one of those non-relatives is more genetically (and hence, phenotypically) similar to the focal individuals’ broodmates than the other. Third, we tested whether males use family-referencing based on the phenotype of the mother by determining whether they discriminate between maternal (i.e. shared mother) and paternal (i.e. shared father) half-siblings, neither of which they developed with. We asked each of these questions in the context of male-male competition because guppies have male-limited color patterns that are highly polymorphic and partially y-linked (Hughes et al. 2013); these patterns provided a reliable phenotypic marker of paternity that we used to infer variation in relatedness among the males derived from split-paternity broods (see below). Fourth, we determined the relative importance of visual and/or olfactory cues in phenotype matching by testing for discrimination between a sibling and a non-relative when only visual, only olfactory, or both visual and olfactory communication was allowed. We asked this question in the context of female mate choice (i.e. inbreeding avoidance), because mate choice interactions are more amenable than male-male competition to the divided tank format that we used to restrict communication modalities.

### Study system and husbandry

Guppies have a promiscuous mating system with internal fertilization (Liley and Stacey 1983; Houde 1997). We used lab-reared descendants of the ‘Houde’ tributary of the Paria river (GPS: PS 896 886). Male color patterns are highly y-linked in this population (Houde 1992; Daniel et al. 2019). This population has very high rates of multiple paternity, with multiple sires contributing to 95% of broods, and mean relatedness among broodmates at 0.36 (compared to 0.5 for full-siblings) (Hain and Neff 2007). Additionally, male guppies frequently mate with multiple females, meaning that individuals often have half-siblings born to different mothers (Johnson et al. 2010). These factors reduce the reliability of a kin template based on the phenotypes of broodmates or the mother. Sires provide no parental care, meaning that is not possible for individuals to base the kin template on the phenotypes of their fathers. Therefore, selection is predicted to favor the evolution of self-referencing rather than family-referencing.

We performed behavioral experiments to test for kin discrimination in the contexts of female inbreeding avoidance, and male avoidance of intrasexual competition with kin. Male reproductive behaviors include following the female, attempting sneak matings (i.e. thrusting the gonopodium at the female’s gonopore without solicitation), and performing courtship displays (Rodd and Sokolowski 1995; Houde 1997). When multiple males simultaneously pursue a female, they frequently compete for physical access to the female’s gonopore. This consists of interruption behavior, wherein the trailing male darts around the leading male, usurping his position behind the female (Gorlick 1976; Bruce and White 1995). Males spend substantial time jockeying for position in this way, and interruptions reduce the copulation opportunities of the interrupted male (Daniel and Williamson 2020). Male guppies spend less time following females that are currently being pursued by kin rivals than females currently being pursued by non-kin rivals, and consequently interrupt kin rivals less than non-kin rivals. This is a form of nepotism that increases the mating opportunities of closely related rivals (Daniel and Williamson 2020). Therefore, we used interruption behaviors and time spent following the same female to test for kin discrimination by males.

We tested for female kin discrimination in divided tanks that allowed us to manipulate whether females could assess the visual and/or olfactory cues of male suitors held in separate compartments. We used the amount of time that females spent associating with each male compartment as a measure of mating preference, and hence kin discrimination. To validate the use of association time as a measure of female mate choice, we re-tested a subset of females with the same suitors but allowed them to freely interact. Association time was strongly correlated with the preference behaviors that females exhibited when freely interacting with males (see Results Expt D): orienting, approaching, and/or moving in front of the male with a “glide” response (Rodd and Sokolowski 1995; Houde 1997). Furthermore, both association time and orienting towards the male have been widely used to measure female preference in this species (Bischoff et al. 1985; Kodric-Brown 1989, 1992; Houde and Torio 1992; Brooks and Caithness 1995; Houde and Hankes 1997; Rosenqvist and Houde 1997; Hibler and Houde 2006; Hampton et al. 2009; Daniel et al. 2019), including in studies of inbreeding avoidance (Daniel and Rodd 2016), and these behaviors predict male mating success (Bischoff et al. 1985; Kodric-Brown 1989).

In our lab population, guppies were held in 54 L stock tanks (60 x 30 x 30 cm) and we moved fish between these tanks every 1 or 2 generations to minimize inbreeding. The fish used in our experiment were held in 5 L tanks (30 x 15 x 20 cm) lined with neutral colored gravel on the bottom and lit by full-spectrum fluorescent bulbs. We maintained a 12:12 h light:dark cycle at 25-26° C, and fed either Tetramin flake fish food or live nauplii larvae of *Artemia salina* twice daily. All individuals were sexually mature (at least 100 days old) before they were used in the experiment.

### General procedures

In this section, we outline procedures common to all four experiments. We generated the fish used in our experiment from a 6-generation breeding design in which all breeding pairs contained individuals that were at most second cousins, to avoid inbreeding. All descriptions of kinship among experimental fish (e.g. full-brothers, non-kin, etc.) are based on relatedness within this breeding design.

Female guppies mate relatively indiscriminately as virgins, but become choosy after an initial mating (Houde 1997; Daniel and Rodd 2016). Therefore, we mated the females used in our behavioral trials to an unrelated male 24 h prior to testing. All trials were performed between 9:30 AM and 12 PM, and behaviors were live-scored manually using JWatcher™, v 1.0 (Blumstein and Daniel 2007). Fish were fed 30 min prior to each trial to discourage foraging behaviors, and females were placed in the observation tank 15 min before the start of the trial to acclimate. When selecting males to use together in a trial, we chose males by eye that were similar in body size and amounts of orange and black coloration on the body and tail, as these traits predict attractiveness in some guppy populations (Houde 1987, 1997). To avoid pseudoreplication, we only used one focal individual from each breeding pair. Except where otherwise indicated, we never used any individual in more than one trial. We had a final sample size of n = 30 trials for each experiment (Expts A-C), and for each modality test (Expt D). We ran all trials for 30 min.

### Experiment A: Do guppies recognize kin through self-referencing?

To determine whether guppies recognize kin through self-referencing, we asked whether males from split-paternity broods discriminate between their full- and maternal half-brothers. Both types of brothers share a mother, develop together *in utero*, and have similar opportunities to associate with one another after birth. Consequently, we reasoned that a kin template based the phenotypes of broodmates or the mother should, on average, be equally similar to the phenotypes of both full- and maternal half-brothers. Therefore, if guppies discriminate between these types of siblings it can only be explained by self-referencing.

To generate the split-paternity broods, we mated virgin females to two males each. We chose two males that differed substantially in color pattern, so that the paternity of sons could be reliably assessed from color pattern (Daniel et al. 2019). These sires were removed from the female’s tank after she became visibly gravid. After giving birth, the mother was removed from the tank within 24 h. Broodmates were reared together in the tank and, as they matured, were sorted by sex into separate tanks to prevent matings. Trials were conducted 4 to 6 weeks after all males in a brood had matured (males matured at roughly 100 days of age). We avoided bias in the proportion of broodmates that were full versus half-siblings in two ways. First, we excluded broods (n = 18) in which there was a ratio greater than 3:2 in favour of either sire’s sons (as identified by color pattern). Second, to choose a focal male from a brood, we randomly selected one of the two sires contributing to that brood, and then randomly selected one focal male from amongst his sons. Thus, a sire’s share of paternity was unrelated to the probability we selected one of his sons as focal.

We placed the focal male in a large observation tank (90 x 45 x 37 cm) along with one full-brother and one half-brother (both from the same brood as the focal male), and 3 mature females unrelated to all 3 males. We recorded the number of times the focal male interrupted each of his rivals, how much time he spent pursuing the same female as each of his rivals, and the proportion of each male’s courtship displays to which the courted female responded positively (by orienting, approaching, and/or “gliding”). Both measures of male-male competition (time spent pursuing the same female, and number of interruptions) have similar biological interpretations and, in all three male-male competition experiments, were significantly positively correlated (all *P* <0.001) and had moderate to high coefficients of determination (*R*^2^ of 0.523 – 0.743; see Supplementary Methods and Supplementary Figure S1). We therefore analyzed only number of interruptions.

### Experiment B: Do guppies recognize kin through family-referencing based on broodmates?

To determine whether guppies use family-referencing based on the phenotypes of broodmates, we asked whether focal males discriminate between two non-kin males, one of which is phenotypically similar to the focal male’s broodmates and the other which is not. Our procedures were similar to those described in experiment A, in which focal males developed *in utero* and after birth with their full- and maternal half-brothers. However, here we tested each focal male with two individuals to which they had not previously been exposed, and to which they were not related. One of these unrelated individuals was a “step-brother” – a male sired by the father of the focal male’s maternal half-siblings (Supplementary Figure S2). Because the focal male developed with his maternal half-siblings, the step-brother was genetically (and hence, phenotypically) similar to some of the focal male’s broodmates. In contrast, the other non-kin male, who was unrelated to the focal male and to all the focal male’s broodmates, should not be phenotypically similar to any of the focal male’s broodmates. Consequently, family-referencing based on broodmates should cause the focal male to incorrectly perceive the “step-brother” to be more closely related to him than the other non-kin male. However, the focal male is not related to either the “step-brother” or the other non-kin male; therefore, if the focal male uses self-referencing, he should not discriminate between them. Similarly, both the “step-brother” and non-kin male are unrelated to the focal male’s mother; therefore, if the focal male uses family-referencing, he should not discriminate between them. Following the protocols outlined for Experiment A, we assayed focal males’ behaviors towards the two males in the behavioral trials.

### Experiment C: Do guppies recognize kin through family-referencing based on the mother?

To test for family-referencing based on the mother’s phenotype, we asked whether males from single-paternity broods discriminate between their maternal half-brothers born in a subsequent brood, and their paternal half-brothers (born to a different dam). Family-referencing based on the mother’s phenotype should lead focal males to perceive maternal half-siblings as more closely related than paternal half-siblings. However, because both types of half-siblings are related to the focal individual at r = 0.5, self-referencing should not lead to discrimination between them. Additionally, the focal individual developed in, and was reared with, a single-paternity brood that did not contain offspring sired by the father of his maternal half-brother; this means that if the focal male uses family-referencing based on broodmates, he should not discriminate between these two types of half-brothers.

We generated single-paternity broods by mating each virgin female to a single male. We selected a focal male from her first brood. We then we re-mated each parent to a different individual to generate maternal and paternal half-brothers of the focal male. We conducted behavioral trials following protocols identical to those used in the experiments above, except that the males in a trial consisted of a focal male, one of his maternal half-brothers, and one of his paternal half-brothers. Because of the time-frame involved in each focal males’ parents re-mating and producing second broods, both maternal and paternal half-brothers were approximately 1 month younger than the focal male, but they were all mature at the time of the trial.

### Experiment D: Do guppies use visual and/or olfactory cues to recognize kin?

To determine the relative importance of visual and olfactory cues in phenotype matching, we assessed female mating preferences for a full-brother versus a non-kin male in a divided tank format that either permitted both communication modalities, or restricted communication to only visual cues or only olfactory cues. Each focal female was derived from a different, singly-mated breeding pair.

To test female preference when both visual and olfactory cues are available, we placed females in the center compartment of a divided tank, with one male placed in each of the two compartments at either end of the tank (Supplementary Figure S3a). Each end compartment was separated from the female by two dividers of clear Plexiglas^®^ perforated with 3 mm diameter holes spaced 1 cm apart, allowing visual and olfactory communication among fish. We used two dividers to separate each compartment for consistency with the other setups (described below). The female’s compartment was delineated into a “neutral” zone in the middle (7 cm wide), and two zones of association (5 cm wide, corresponding roughly to 3 body lengths), each adjacent to an end compartment. As a measure of female preference, we recorded the time the female spent with her snout in each zone of association. Males were added to the end compartments, and the trial began 15 minutes later when the female was added. Each trial ran for 15 min.

To test for the role of visual cues alone in kin recognition, we observed females using the same protocol and in a divided tank identical to the one described above, except that the outer divider separating each end compartment was not perforated and was thus watertight (Supplementary Figure S3b). This prevented olfactory cues from moving between compartments.

Finally, to assess the role of olfactory cues alone in kin recognition, we tested females in a setup identical to the previous description, except that males were not present in the observation tank. Instead, males were each held in a separate tank containing 1 L of water, out of view of the female. Silicon tubing (3 mm diameter) slowly siphoned water from each male’s tank (following McLennan and Ryan 1997) and released it near the bottom of the space between the pairs of internal dividers in the female’s tank. This allowed olfactory cues to move through the perforated divider and reach the female (Supplementary Figure S3c). Valves attached to the tubing held the flow rate constant at 2 mL/min throughout each trial. Prior to experimentation, testing with dyes indicated that our setup produced a marked gradient of water flowing from each side, meeting at the center of the female’s compartment. This gradient was stable over the timeframe of our trials. To ensure that olfactory cues were at high enough concentrations to be detected, the males were held in their tanks for 24 h prior to the trial, without food. Non-experimental females were held in tanks adjacent to the males’ tanks to provide visual stimulation. Both water sources were running when the focal female was placed in the observation tank.

Across all three setups, for each trial, we randomized which side held (or received water from the tank of) the full-brother or the non-kin male. Between each trial, we cleaned the observation tank (and male holding tanks and silicon tubing, when applicable) by flushing with soapy water, hydrogen peroxide, and then thoroughly rinsing (following McLennan and Ryan 1997; Makowicz et al. 2016). Across all three setups, we excluded 11 trials in which the female did not approach both zones of association within the first 4 minutes (and therefore may have had limited information about her options for much of the trial). To validate association time as a measure of female preference, we immediately re-tested a subset (n = 15) of the females used in the visual and olfactory communication trials with the same two males in open association trials. We followed the same protocols as for Expts A, B and C, including scoring the proportion of male courtship displays towards which females responded positively. We asked whether female association time was correlated with the preference behaviors that females exhibited when allowed to freely interact with males.

### Statistical Analyses

We performed all analyses in R, v 3.5.3. All analyses were two-tailed. For Expts A, B and C, we asked whether focal males directed different numbers of interruptions towards different types of stimulus males. For experiment D, we asked whether females associated more with one type of male than another.

The behaviors that a focal individual directs towards the two stimulus males in a given trial are not independent of one another. This non-independence arises, in part, because of an inevitable trade-off between the time and energy that a focal individual expends directing behaviors (i.e. interruptions, or association time) towards one stimulus male versus another. To accommodate this non-independence, we calculated each focal individual’s behaviors towards one type of stimulus male (i.e. number of interruptions towards the full-brother, step-brother, or maternal half-sibling for Expts A, B, and C; association time with the brother for Expt D) as a proportion of the total behaviors that the focal individual directed towards both types of stimulus males–e.g. for Expt A: number of interruptions towards the full-brother / (number of interruptions towards the full-brother + non-kin male). Proportions are often non-normal if sample sizes are low, and if there is a large number of extreme (close to 0 or 1) values (Warton and Hui 2011). For all experiments, our proportion data did not differ significantly from normality (Shapiro-Wilk test, package *stats* v 3.5.3, all *P* > 0.05), likely because sample sizes were moderate (n = 30) and there were few extreme values. For this reason, we did not apply transformations. To test for kin discrimination, we asked whether these proportions differed significantly from 0.5 (the null expectation in the absence of kin discrimination), using one-sample t-tests (package *stats* v 3.5.3). For Expt D, we also asked whether female association bias differed depending on which cue modality(ies) were available. We compared female association bias between each pair of stimulus modality setups using two-sample, unpaired t-tests with equal variances.

For Expts A, B, and C, we wanted to determine whether the focal male’s behavior could have been altered by females responding differently to the focal male depending on which stimulus male he was competing with. We therefore asked whether the responsiveness of females (measured as the proportion of the focal male’s courtship displays to which females responded positively) differed between portions of the trial in which he was pursuing the same female as one type of stimulus male versus the other type of stimulus male. Female responsiveness had a zero-inflated distribution and was non-normal even after applying transformations. Consequently, we used paired, two-sample Wilcoxon signed-rank tests (package *coin*, v 1.3-1), and calculated the exact test, to compare female responsiveness to the focal male between the two different competitive contexts.

For Expt D setups that permitted visual communication, we wanted to know whether full-brother and non-kin stimulus males differed in their courtship effort, as this could have biased female association behavior. We compared the number of courtship displays performed by the full-brothers and non-kin males. We performed this test separately for the visual and olfactory, and olfactory only, setups. To accommodate non-normality that persisted despite transformations, we used two-sample, paired Wilcoxon signed-rank tests, and calculated the exact test.

We also wanted to know whether association bias predicts the preference behaviors females exhibit when freely interacting with males. We asked whether the proportion of association time that females spent with the full-brother was correlated with the difference in female responsiveness towards the full-brother’s courtship and the non-kin male’s courtship. Association bias and difference in responsiveness met parametric assumptions (homoscedastic residuals, and normality – multivariate Shapiro-Wilk test (package *RVAideMemoire* v 0.9-73): *P* = 0.827). Therefore, we used Pearson’s product-moment correlation to ask whether these two measures of female preference were correlated.

## Results

### Experiment A

As a test of self-referencing, we asked whether males discriminate between their full- and maternal half-brothers. The proportion of interruptions that focal males directed towards their full-brothers rather than their half-brothers was significantly lower than expected in the absence of kin discrimination (mean ± SE = 0.385 ± 0.039, *t_29_* = −2.925, *P* = 0.007; Figure 1). The proportion of male displays to which females responded positively was not significantly affected by whether she was simultaneously pursued by the focal male and his full-brother versus the focal male and his half-brother (full-brothers: 0.203 ± 0.032, half-brothers: 0.201 ± 0.025, *Z* = −0.142, *P* = 0.891). Thus, the bias in focal male competitive behavior cannot be explained by changes in female responsiveness associated with between-male relatedness.

**Figure 1:**
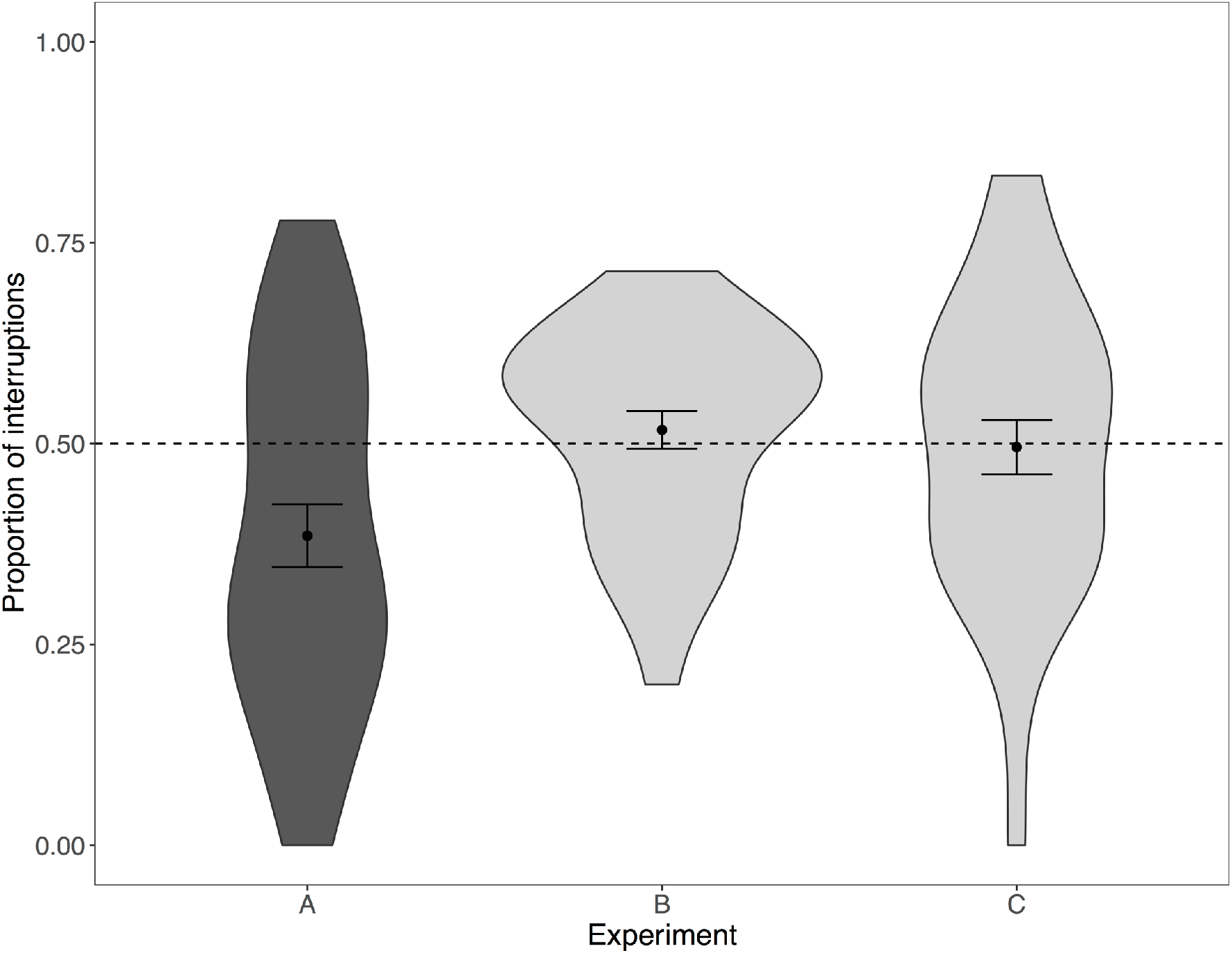
The proportion of the focal male’s interruptions that were directed towards the full-brother rather than the half-brother (Expt A), the “step-brother” rather than other non-kin male (Expt B), or the maternal half-brother rather than the paternal half-brother (Expt C). Violins depict the distribution (kernel density) of the data. The black dot represents the mean ± standard error. Dark grey violins denote significant kin discrimination; light grey denotes no significant kin discrimination. The dashed line indicates the proportion expected in the absence of kin discrimination (0.5).

### Experiment B

To test for family-referencing using the phenotypes of broodmates, we asked whether males discriminate between their unrelated “step-brothers” and other non-kin males. The proportion of interruptions males directed towards their “step-brothers” rather than other non-kin males was not significantly different from the null expectation in the absence of kin discrimination (0.517 ± 0.025, *t_29_* = 0.721, *P* = 0.477; Figure 1). Female preference behavior was not significantly affected by whether the female was interacting with the focal male and his “step-brother” or the focal male and the other non-kin male (“step-brother”: 0.210 ± 0.028, other non-kin: 0.219 ± 0.026, *Z* = −0.559, *P* = 0.582).

### Experiment C

To test for family-referencing using the mother’s phenotype, we asked whether males discriminate between their paternal and maternal half-brothers. The proportion of interruptions that focal males directed towards their maternal half-brothers rather than their paternal half-brothers was not significantly different from the null expectation in the absence of kin discrimination (0.496 ± 0.034, *t_29_* = −0.130, *P* = 0.898; Figure 1). Female preference behavior was not significantly affected by whether the female was being pursued by the focal male and the maternal half-sibling, or the focal male and the paternal half-sibling (maternal: 0.221 ± 0.021, paternal: 0.199 ± 0.026, *Z* = 0.651, *P* = 0.52).

### Experiment D

To test for the role of visual and olfactory cues in kin recognition, we asked whether females associate preferentially with non-kin males over their full-brothers in divided tank setups that manipulated which cue modalities females could assess. As predicted, when both visual and olfactory cues were available, females spent significantly less than half of their total association time associating with their full-brother (0.416 ± 0.029, *t_29_* = −2.89, *P* = 0.007; figure 2). When only visual cues were available, females did not exhibit an association bias (full-brother: 0.517 ± 0.027, *t_29_* = 0.611, *P* = 0.546; figure 2). Full-brothers and non-kin males performed similar numbers of courtship displays, both in the visual and olfactory trials (full-brother: 3.33 ± 0.35, half-brother: 3.5 ± 0.298, *Z* = −0.816, *P* = 0.419), and the visual only trials (full-brother: 2.2 ± 0.269, half-brother: 2 ± 0.173, *Z* = 0.372, *P* = 0.713). Thus, female association preferences cannot be explained by differences in male courtship effort. When only olfactory cues were available, females associated less with their full-brothers than expected in the absence of kin discrimination (full-brother: 0.430 ± 0.023, *t_29_* = −3.01, *P* = 0.005; figure 2). Female association bias was strongly, positively correlated with the difference in the proportion of each male’s courtship displays to which the female responded positively (df = 13, *R*^2^ = 0.621, *P* < 0.001; Supplementary Figure S4). Association bias therefore reflects the preferences that females exhibit when allowed to freely interact with males.

**Figure 2:**
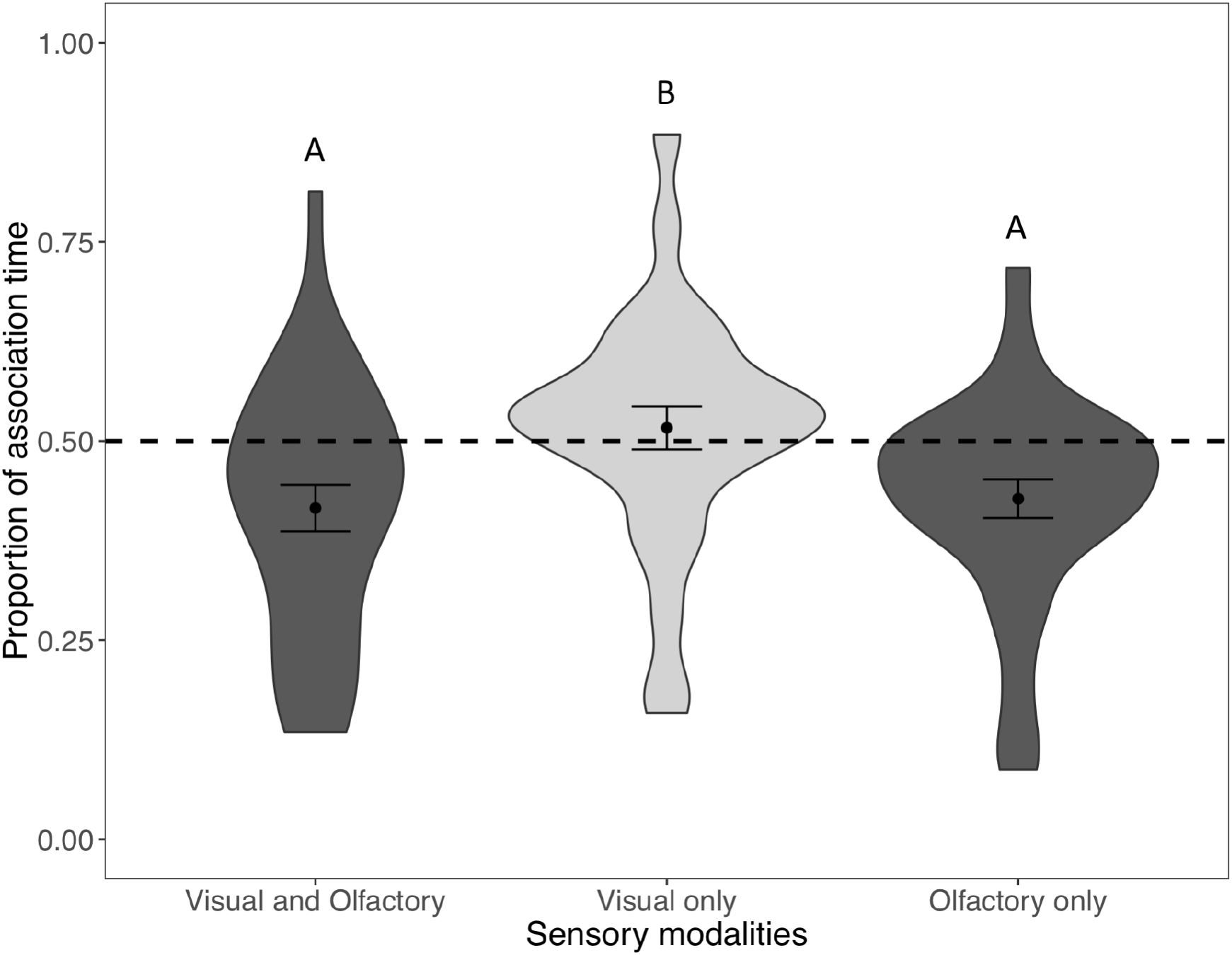
The proportion of total association time that the female spent near her full-brother rather than their half-brother, when both visual and olfactory, only visual, or only olfactory cues were available. Violins depict the distribution (kernel density) of the data. The black dots represent the means ± standard errors. The dashed line indicates the proportion expected in the absence of kin discrimination (0.5). Dark gray indicates significant kin discrimination; light gray indicates no significant kin discrimination. Same letters indicate no significant difference in means among sensory modalities; different letters indicate significantly different means among sensory modalities.

We next asked how female association bias differed depending on which cue modalities were available. Females showed a stronger association bias favoring non-kin when both visual and olfactory cues were available than when only visual cues were available (*t_58_* = −2.529, *P* = 0.014). Female preference for associating with non-kin males was not affected by whether visual and olfactory cues, or only olfactory cues, were available (*t_58_* = −0.377, *P* = 0.708). Females showed a stronger association bias favoring non-kin when only olfactory cues were available than when only visual cues where available (*t_58_* = 2.423, *P* = 0.019).

## Discussion

Our results provide insight into the proximate mechanisms underpinning kin recognition. We found that male guppies discriminated between full- and half-brothers, despite sharing a mother and developing *in utero* with both types of brothers. Therefore, family-referencing based on the mother’s phenotype or the phenotype of broodmates cannot explain this discrimination amongst full- and half-brothers. In contrast, self-referencing relies on the evaluator’s own phenotypic similarity to conspecifics, and can thus explain this discrimination between full and maternal half-siblings. These results provide the first conclusive demonstration, to our knowledge, of self-referencing in a live-bearing species.

We found no evidence of family-referencing. Focal males did not discriminate between two types of non-kin individuals, one of which–a “step brother”–was similar to their broodmates. This result suggests that guppies do not use phenotypic information about their broodmates to form a kin template. Second, males did not discriminate between unfamiliar maternal and paternal half-siblings, suggesting that guppies do not use the phenotype of their mother to form the kin template. This lack of evidence for family-referencing based on broodmates or on the dam cannot be attributed to lack of statistical power because sample sizes were consistent across experiments, and within-group variance in behavior was greatest in the test of self-referencing, for which we found significant evidence of kin discrimination. Furthermore, the lack of evidence of family-referencing is unlikely to have resulted from guppies being unable to discriminate between the levels of phenotypic and genetic variation presented in our experiments (i.e. variation between maternal and paternal half-siblings, and between a “step-brother” and another non-relative), since we found that guppies readily distinguished between full- and half-siblings. Collectively, our results indicate that phenotype matching utilizes a kin template based on self-assessment, and that exposure to the cues of kin *in utero* or after birth is unimportant for the ontogeny of kin recognition in this species.

That guppies rely on self-rather than family-referencing supports the prediction that self-referencing is more likely to evolve than family-referencing in systems with high levels of multiple paternity, like those documented in our study population (Hain and Neff 2007). Similarly, a previous study has shown that differences in the level of promiscuity between alternative life histories predicts the use of self-referencing in bluegill sunfish (*Lepomis macrochirus*) (Hain and Neff 2006). Sunfish have external fertilization, which allowed the authors to manipulate potential exposure to kin from conception onwards using *in vitro* fertilization. The set of breeding crosses we used to disentangle self- and family-referencing can be applied to live-bearing species, expanding the range of study systems in which researchers can disentangle these kin recognition systems.

We also investigated the importance of visual and/or olfactory phenotypes in kin recognition. Consistent with previous work (Daniel and Rodd 2016), female guppies discriminated between full-brothers and non-kin males when both visual and olfactory cues were available. Females discriminated to a similar degree when only olfactory cues were available, but did not discriminate when only visual cues were available (despite equivalent sample sizes and similar variances). Taken together, these results indicate that olfactory cues are both necessary and sufficient for kin recognition, while visual cues are unimportant for kin recognition. Reliance on olfactory phenotypes for kin recognition is perhaps unsurprising considering the use of self-rather than family-referencing to form the kin template; olfactory cues should be more amenable than visual cues to self-inspection in the guppy. In combination with several previous studies that have also documented olfactory-based kin recognition in teleost fishes (Mann et al. 2003; Mehlis et al. 2008; Makowicz et al. 2016), our results suggest that olfactory cues may be a common cue for fish kin recognition.

Our results indicate that phenotype matching depends on olfactory cues, yet the specific odor components involved remain unclear. In many vertebrate taxa, genotypic similarity at the major histocompatibility complex (MHC) is associated with differences in mate choice and social behavior (Bonneaud et al. 2006; Schwensow et al. 2008; Evans et al. 2012; Rymešová et al. 2017; Winternitz et al. 2017), suggesting a possible role in kin recognition. MHC genotype can influence individual odor because these proteins are present in saliva, urine, and sweat where they can be assessed by the olfactory system, and because their role in immunity can shape gut microbiome composition (Kubinak et al. 2015). These genes encode transmembrane proteins important for self-recognition as part of immune function because they bind specific self- and pathogen-derived antigens, which they present to the immune system. MHC genes are extremely polymorphic, likely because of balancing selection exerted by co-evolution with pathogens, and sexual selection (e.g. Bernatchez and Landry 2003; Solberg et al. 2008; Winternitz et al. 2017). For example, female mate choice in the three-spined stickleback, *Gasterosteus aculeatus*, appears to involve females choosing males whose MHC genotypes compliment their own, such that their offspring will have optimal MHC diversity (Reusch et al. 2001; Aeschlimann et al. 2003; Wegner et al. 2003, 2008; Milinski et al. 2005; Eizaguirre et al. 2009; Kalbe et al. 2009; Lenz et al. 2009). Because MHC genes are highly polymorphic, odor cues associated with MHC genotype likely provide a reliable signal of relatedness. Mate choice for optimal MHC diversity may, therefore, also function in inbreeding (and outbreeding) avoidance. In guppies, inbred individuals suffer higher loads of the parasite *Gyrodactylus turnbulli*, and are slower to clear infections than outbred individuals (Smallbone et al. 2016). Reduced MHC diversity likely contributes to these deleterious effects of inbreeding. The potential role of MHC-dependent odor cues in shaping kin recognition in the context of male-male competition has not, to our knowledge, been investigated, and is fruitful avenue for future work.

Despite our finding that females do not use visual cues for kin recognition, much previous work has demonstrated that male visual ornaments play an important role in guppy mate choice. For example, females from many populations prefer males with larger and more saturated orange spots (e.g. Houde 1997), which likely serve as honest signals of male quality (Nicoletto 1993; Grether 2010). Combined with this literature, our findings highlight that female mate choice is multi-modal (Guilford and Dawkins 1993; Rowe and Guilford 1999) in the guppy. Females may use different cue modalities to assess different aspects of mate suitability. For example, female sticklebacks select males based jointly on olfactory cues that indicate MHC genotype complementarity, and on the intensity of coloration, which indicates whether a male has the specific MHC alleles that provide resistance against current diseases (Milinski 2014). The potential interplay between female assessment of mate quality through visual cues and mate compatibility (including relatedness) through olfactory cues presents an interesting avenue for further inquiry.

As in our behavioral trials, guppies often make mate choice and competitive decisions while multiple conspecifics are in the vicinity. This raises the question of how guppies may be able to ascertain which of multiple conspecifics are the source of a given olfactory cue or profile. Interaction between the olfactory and vibration-detection (i.e. lateral line) sensory systems may allow guppies to attribute scents to particular individuals. For example, sharks use their lateral line for “eddy” chemotaxis (Atema 1995, 1996, 1998), detecting small-scale eddies produced by movement to locate odor-producing objects (Gardiner and Atema 2007). Similar interactions between olfactory and movement cues (e.g. the vibrations the male guppies perform when courting (Houde 1997), or the movement of a stimulus male pursuing a female) may help guppies to determine which individual suitor, or rival is the source of a given scent. Our future work will ask whether courtship vibrations help females assess relatedness to their suitors’ (and whether males adjust this behavior strategically depending on their relatedness to the female).

That guppies use both self-referencing and olfactory phenotypes for kin recognition raises several questions about the mechanisms of kin template formation. Despite ample exposure to kin (and their odor cues) during early life, our results suggest that individuals incorporate only their own phenotypes into the kin template. How, then, do individuals discriminate between their own olfactory cues and those of conspecifics to form the template? Possible explanations include differences in the salience of self-versus non-self cues, which may occur at different local concentrations in the water immediately surrounding an individual, or differences in sensitivity to those cues. Differences in olfactory sensitivity to self-versus non-self cues could involve a genetic component similar to the innate immune system, in which the binding or rejection of olfactory ligands is determined by their similarity to MHC-encoded surface proteins (Gerlach et al. 2008).

Another intriguing question concerns the timeframe over which the kin template is formed and retained. For species using family-referencing, a reliable kin template must be formed during early life or some other developmental window within which exposure is limited to ones’ close kin, and then stored for later use. In contrast, for self-referencing, the kin template could be formed at any period throughout development, and could be updated over the life course through continuous self-assessment (e.g. Lizé et al. 2009). The ability to update the kin template should be advantageous if templates decay over time (e.g. as memory fades). Furthermore, for taxa other than the guppy that use kin recognition systems only intermittently (e.g. because of defined breeding seasons), energetic costs could be reduced by letting the kin template decay and subsequently replacing it. Indeed, seasonal plasticity of neurological substrates is believed to reduce the metabolic costs of learned behaviors in other taxa (e.g. Mateo 2010). Exploring whether and how the kin template is either maintained or reformed over the life course could provide insight into the neurological processes of phenotype matching, and the selective forces favoring self-versus family-referencing.

### Conclusions

The design of our experiments allowed us manipulate potential exposure to the cues of kin from conception onwards, while preserving natural development. Unlike many previous tests of phenotype matching, we were thus able to disentangle self-referencing from family-referencing occurring *in utero* or shortly after birth as explanations for the formation of the kin template that guppies use to recognize kin. Our results demonstrate that, in the guppy, phenotype matching (i) relies on self-referencing, not family-referencing, and (ii) utilizes olfactory rather than visual cues. This finding is consistent with the prediction that self-referencing should be favored in promiscuous mating systems like that of the guppy. Our results resolve key components of the proximate mechanisms underpinning phenotype matching, clarifying the means by which females avoid inbreeding and male engage in nepotism during male-male competition. These findings lay the groundwork for inquiries into the genetic and molecular mechanisms of kin recognition, and their role in the evolution of social behavior.

