## Supplementary Materials for "Kin recognition in guppies uses self-referencing on olfactory cues"

**Supplementary Material**

Supplementary Methods

We used correlation analysis to ask whether the two behavioral measures of male-male competition (time that the focal male spent pursuing the same female as a given stimulus male, and number of times the focal male interrupted a given stimulus male) provided similar information about kin discrimination. Both of these behavioral measures were calculated as proportions (i.e. amount of behavior towards one stimulus male, divided by total response towards both stimulus males). For all three male-male competition experiments, both behavioral measures had a small number of extreme values, and did not deviate significantly from univariate or bivariate normality (Shapiro-Wilk test: all $P > 0.09$; multivariate Shapiro-Wilk test: all $P > 0.57$). For this reason, we did not apply transformations. Additionally, visual inspection revealed a linear relationship between the two behavioral measures for all three experiments (Supplementary Figure S1). We therefore calculated the Pearson product-moment correlation (package *stats*, v 3.5.3).

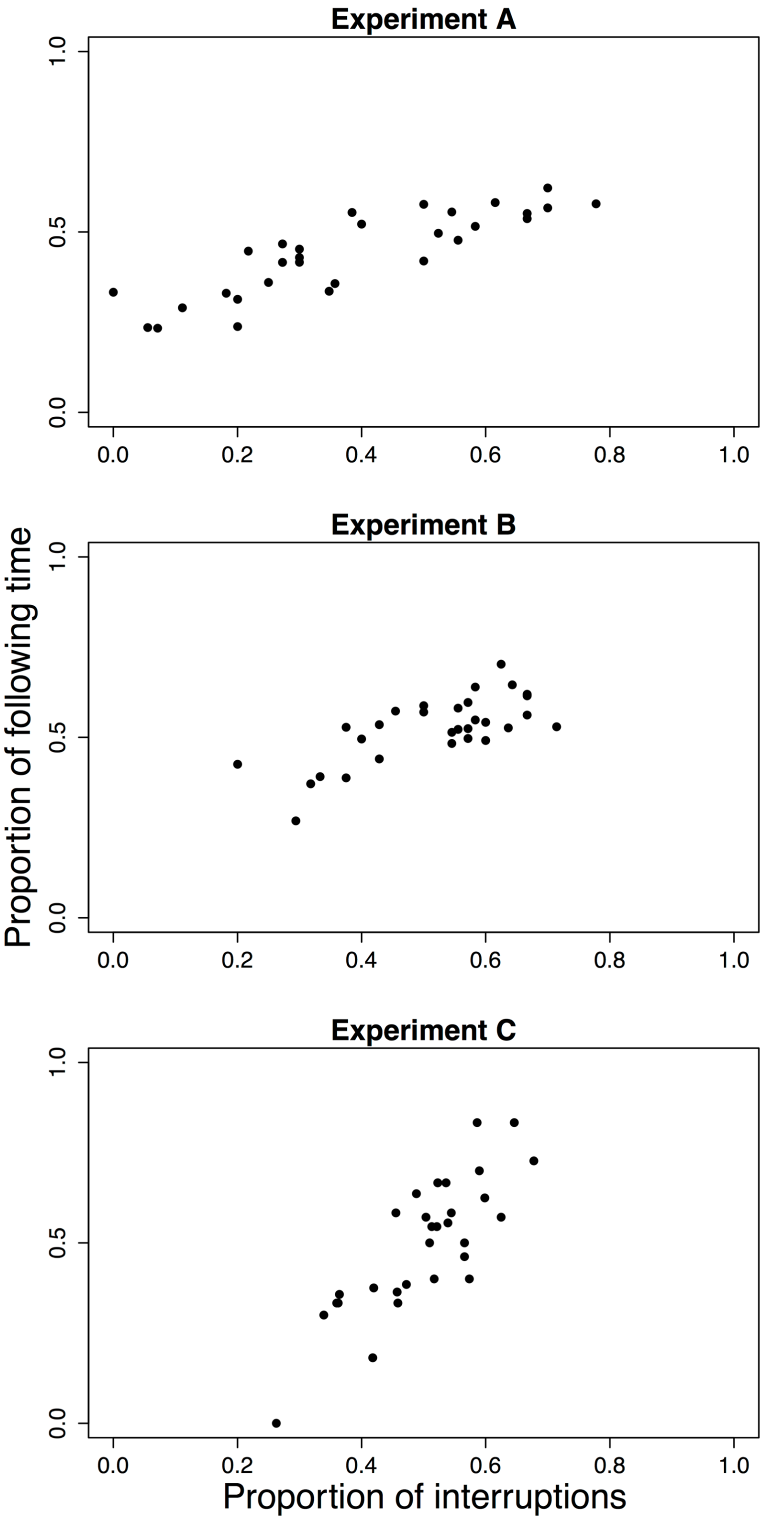


Figure S1. Scatterplots, for Expts A-C, of the two behavioral measures of male-male competition: interruptions, and amount of time spent following the same female.

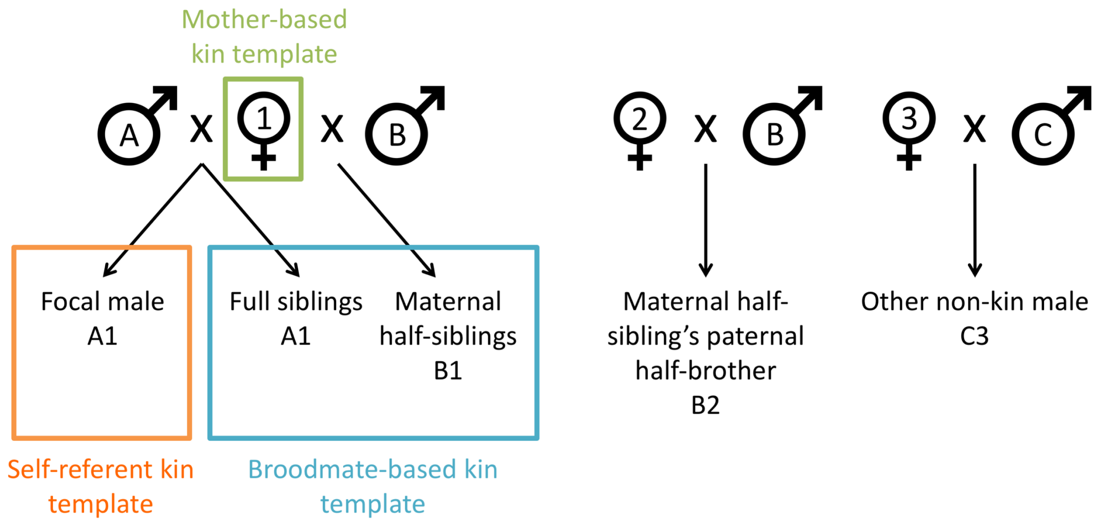


Figure S2. Breeding crosses used to test for family-referencing based on the phenotypes of broodmates (Expt B). Letters correspond to paternal line; numbers correspond to maternal line. Categories of relatedness (e.g. "full siblings") are from the perspective of the focal male. Each focal male (A1) developed in a split-paternity brood, alongside his full-siblings (also A1) and maternal half-siblings (A2). Under self-referencing (orange), the focal male's kin template is based on his own phenotype and reflects phenotypes associated with parents A and 1. Family-referencing on broodmates (cyan) should lead the focal male to incorporate the phenotypes of both full- and maternal half-siblings into his kin template, such that it will reflect the phenotypes associated with parents A, B and 1. Family-referencing on the mother (green) should result in a kin template reflecting parent 1. To test for family-referencing based on broodmates we assayed interactions between a focal male and two unrelated males: a "step-brother" (i.e. a paternal half-brother of the focal individual's maternal half-siblings; B2) and another non-kin male (C3). Since these males do not share a parent with the focal male, neither should be more similar than the other to a kin template based on self-referencing or family-referencing on the mother's phenotype. However, the "step-brother" shares a father with some of the focal male's broodmates (his maternal half-siblings), and should therefore be more similar to a kin template based on broodmates than the other non-kin male (who does not share any parents with the focal male or his broodmates). Therefore, only family-referencing on broodmates should lead to discrimination between the "step-brother" and the other non-kin male.

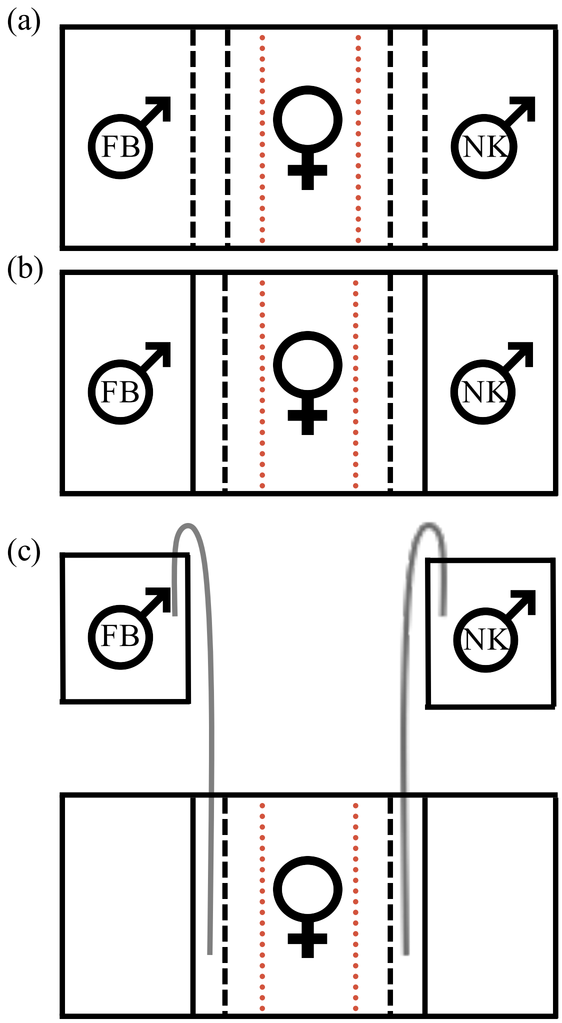


Figure S3. Side views of divided tank setups used to test kin discrimination when (a) visual and olfactory, (b) only visual, or (c) only olfactory cues are available. In (a) and (b), stimulus males (FB = full-brother; NK = non-kin) were randomly assigned to either end compartment, separated from the center compartment containing the focal female by two layers of Plexiglas® (internal black lines). In (a), both layers were perforated to permit olfactory cues. In (b) and (c), the inner layer was perforated while the outer layer was water tight, allowing only visual communication. The female's center compartment was delineated (depicted in orange) into a middle neutral zone and two zones of association, each adjacent to an end compartment. In (c), males were not held in the end compartments, but were instead placed in separate tanks above the divided tank. Silicon tubing (gray) slowly siphoned water from each male's tank into the space between Plexiglas barriers in the divided tank; from there, it flowed towards the female.

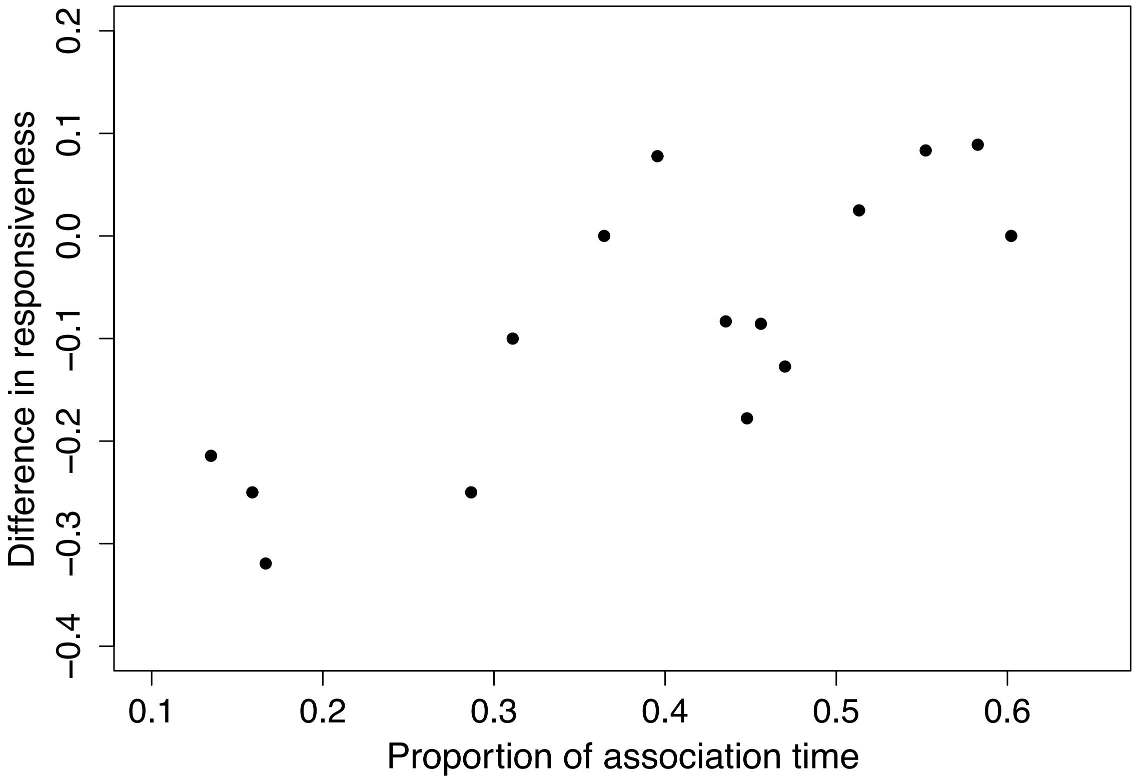


Figure S4. Scatterplot of association bias (proportion of total association time spent associating with the full-brother) and difference in responsiveness (proportion of the full-brother's courtship displays to which the female responded positively, minus the proportion of non-kin male's courtship displays to which the female responded positively).
